# The regulatable *MAL32* promoter in *S. cerevisiae*: characteristics and tools to facilitate its use

**DOI:** 10.1101/061127

**Authors:** Matthias Meurer, Veronika Chevyreva, Bram Cerulus, Michael Knop

## Abstract

Here we describe a set of tools to facilitate the use of maltose and the *MAL32* promoter for regulated gene expression in yeast, alone or in combination with the *GAL1* promoter. Using fluorescent protein reporters we find that under non-inducing conditions the *MAL32* promoter exhibits a low basal level of expression, similar to the *GAL1* promoter, and that both promoters can be induced independently of each other using the respective sugars, maltose and galactose. While their repression upon glucose addition is immediate and complete, we found that the *MAL32* and *GAL1* promoter each exhibit distinct induction kinetics. A set of plasmids is available to facilitate the application of the *MAL32* promoter for chromosomal modifications using PCR targeting and for plasmid based gene expression.

## Introduction

In yeast, promoters that can be regulated – induced or repressed – as a function of conditions or via the addition of compounds have been established as powerful tools for research or biotechnological purposes. Several carbon source dependent promoters (Weinhandl *et al.* 2014) exhibit a high level of dependence on the composition of the growth medium, for example the sugar content. In particular, the *GAL1* promoter (Finley *et al.* 2002) is strongly influenced by the presence of galactose (inducing) or glucose (repressing) in the medium. Induction of the promoter upon addition of galactose to cells grown on raffinose is fast and can be rapidly halted by the subsequent addition of glucose. This feature of the *GAL1* promoter, induction and repression simply based on the addition of compounds, is relatively unique. Other regulatable promoters, such as the heterologous *tet*-promoter, can be regulated only in one direction, e.g. either induced or repressed upon the addition of a compound (Dingermann *et al.* 1992; Gossen and Bujard 1992). In order to reverse the regulation the stimulus needs to be removed, for example, by washing the cells with fresh medium, which is much less convenient. Thus, the *GAL1* promoter (subsequently called *GAL1^pr^)* is the promoter of choice whenever short expression pulses are needed to study a specific process, such as when performing a functional analysis of cell cycle regulation. However, for a number of applications it would be useful to have an additional promoter that can be regulated in a similar manner.

The promoters of the maltose inducible and glucose repressible *MAL* genes seemed to be promising candidates (Weinhandl *et al.* 2014). For example the *MAL62* promoter can be strongly induced by maltose comparable to the *GAL1^pr^*, but under non-inducing or repressing conditions the background expression from the *MAL62* promoter was much higher compared to the *GAL1^pr^* (Levine, Tanouye and Michels 1992; Finley *et al.* 2002) which decreases its usability. Therefore we investigated the regulation of other maltose inducible and glucose repressible *MAL*-promoters in direct comparison with *GAL1^pr^*. Depending on the yeast strain, the fermentation of maltose is governed by up to five unlinked but similar loci, each consisting of 3 genes (Barnett 1976; Carlson 1987). Each locus contains an activator gene, a maltose permease and a maltase. The genes in the different loci are termed *MALxy*, and the nomenclature is such that the first digit denotes the locus (1, 2, 3, 4, or 6) whereas the 2^nd^ digit denotes one of the three genes: *MALx1 for* maltose permease, *MALx2 for* maltase and *MALx3 for* the activator gene (Fig. 1) (Weinhandl *et al.* 2014). Regulated gene expression as a function of the addition of maltose has been well studied and involves the induction of the maltose permease and the maltase encoding genes from a single bidirectional promoter present in the intergenic region of these two genes (Needleman *et al.* 1984; Bell *et al.* 1995). The *MALx3* gene upstream of *MALx1* codes for a transcriptional activator, that regulates the expression of the bi-directional promoter (Chang *et al.* 1988).It is important to note that different yeast laboratory strains contain different numbers of *MAL* loci, but often none are functional for growth on maltose, rendering these strains unable to use maltose as a carbon source.Here we investigate the regulation of maltase and maltose permease gene promoters in direct comparison with *GAL1^pr^*. We then focus on the *MAL32* promoter (subsequently called *MAL32^pr^*) and we outline how this promoter can be used, alone and in combination with *GAL1^pr^*.

**Figure 1.**
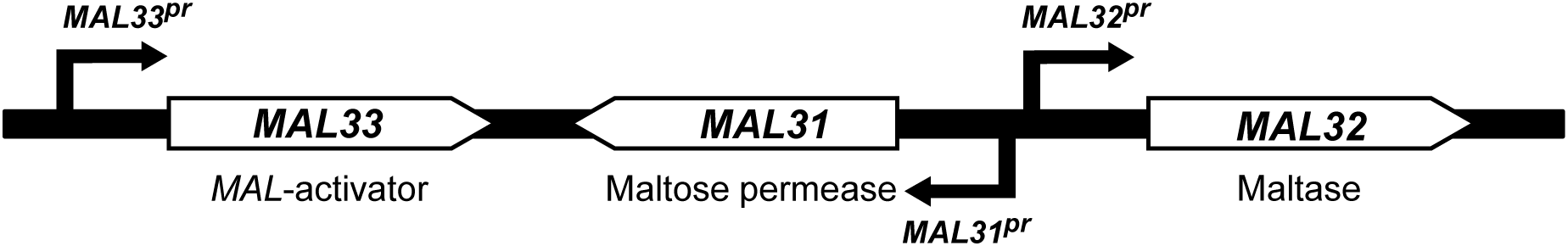
Typical organization of a *MAL* locus, using the *MAL3* locus as an example. For explanations see text.

## Results and Discussion

We used the S288c derived strain ESM356-1 (Knop *et al.* 2005), which is a spore from the diploid strain FY1679 (Winston, Dollard and Ricupero-Hovasse 1995; Wach *et al.* 1997). S288c strains contain two *MAL* loci, *MAL1* and *MAL3*, both containing non-functional activator genes (*mal13* and *mal33*). We decided to investigate all promoters of these loci for regulation by maltose and other carbon sources. We constructed reporter strains expressing *sfGFP*, a bright and fast maturing *GFP* variant (Pédelacq *et al.* 2006), under the control of all promoters from the two *MAL*-loci and compared it to a *GAL1* promoter reporter strain. We tested for expression of the reporter using growth on agar plates and colony pinning using a pinning robot. Promoter activity was quantified in n ≥ 20 colonies per strain using whole colony fluorescence measurements with the help of a fluorescent plate reader. In the presence of the *MAL63* activator (on plasmid pRS415- *MAL63* containing a functional *MAL* activator gene originating from the yeast strain RM11), specific induction of the promoter regulating the maltase and the maltose permease genes was observed (Fig. 2A, B). The achieved expression levels were in the range of 15 – 45% of those observed for *GAL1^pr^*. No induction of the activator (*MAL13^pr^* and *MAL33^pr^*) was observed. We found that *MAL11^pr^* and *MAL12^pr^* showed the highest expression levels when induced by maltose but also higher basal levels compared to *MAL31^pr^* and *MAL32^pr^* when repressed by glucose or uninduced on galactose/ raffinose. *MAL31^pr^* and *MAL32^pr^* both showed low expression levels when repressed or uninduced, however, *MAL32^pr^* exhibited two fold higher levels than *MAL31^pr^* when induced by maltose. Based on these results we decided to use *MAL32^p^* for further work.

**Figure 2.**
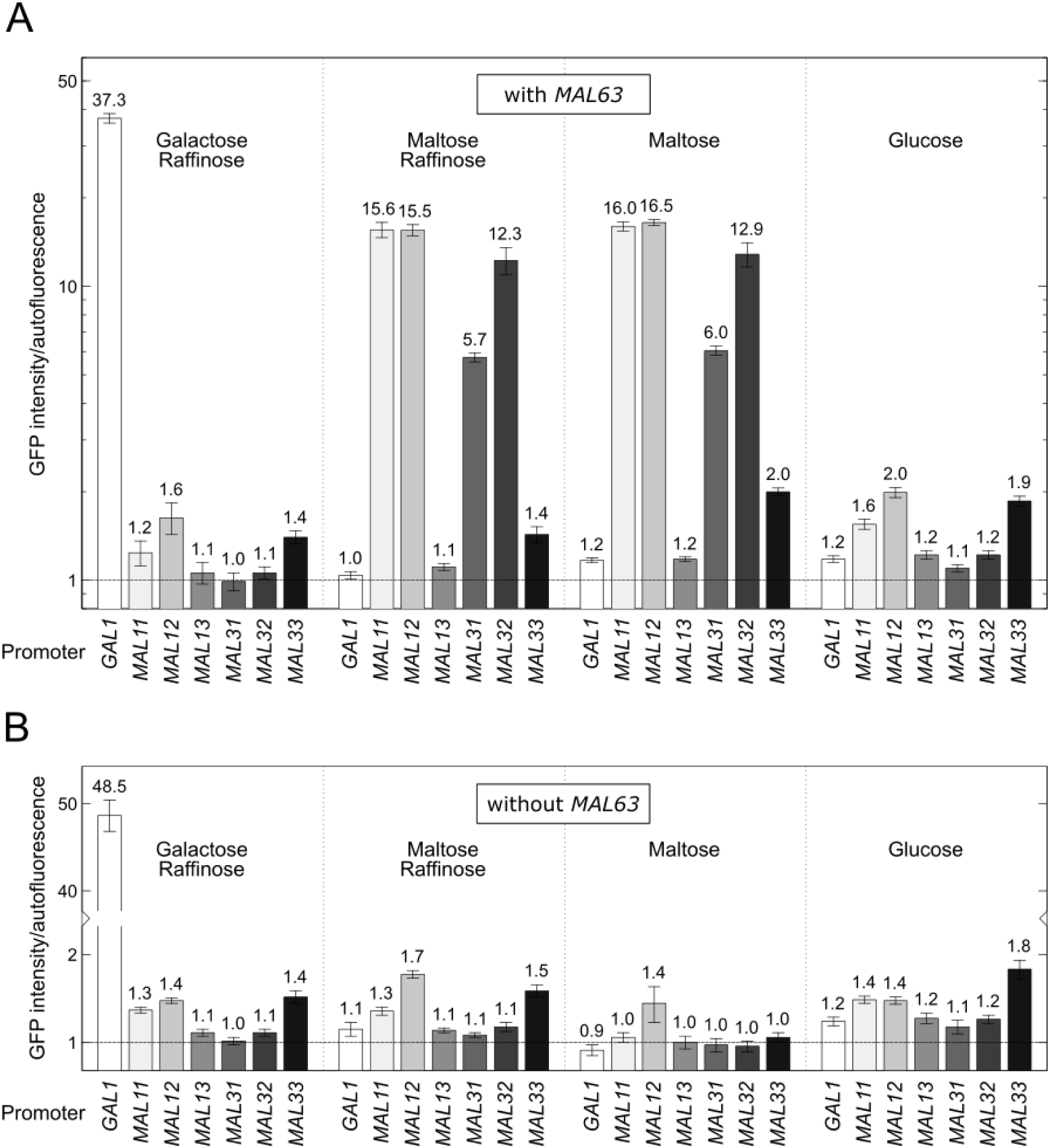
Promoter activity of different *MAL* genes. A genomic promoter duplication strategy (Huber *et al.* 2014) and *sfGFP* as a reporter were used to measure the promoter strength of the indicated *MAL*-genes in the presence (pRS415-*MAL63*) (**A**) or absence (pRS415) (**B**) of a functional *MAL63* activator gene. For strain construction, please refer to Materials and Methods. Fluorescence intensities were determined in colonies (mean ± s.d., *n* ≥ 20 colonies per construct) grown for 24 hours on synthetic complete media lacking leucine containing different carbon sources (2% w/v), as indicated. Intensities were normalized to the background fluorescence of a wild type control strain.

Next, we tested for repression of the promoter using cells grown in liquid medium and a plasmid containing GFP fusions to *MAL32^pr^* or *GAL1^pr^*. This revealed comparable expression levels of both promoters on the respective carbon source (maltose or galactose) and full repression of both promoters in the presence of glucose (Fig. 3). We also tested a truncated *MAL31^pr^* variant (*MAL31^pr^*-short (Levine, Tanouye and Michels 1992)) and found properties comparable to the full length *MAL32^pr^*.

**Figure 3.**
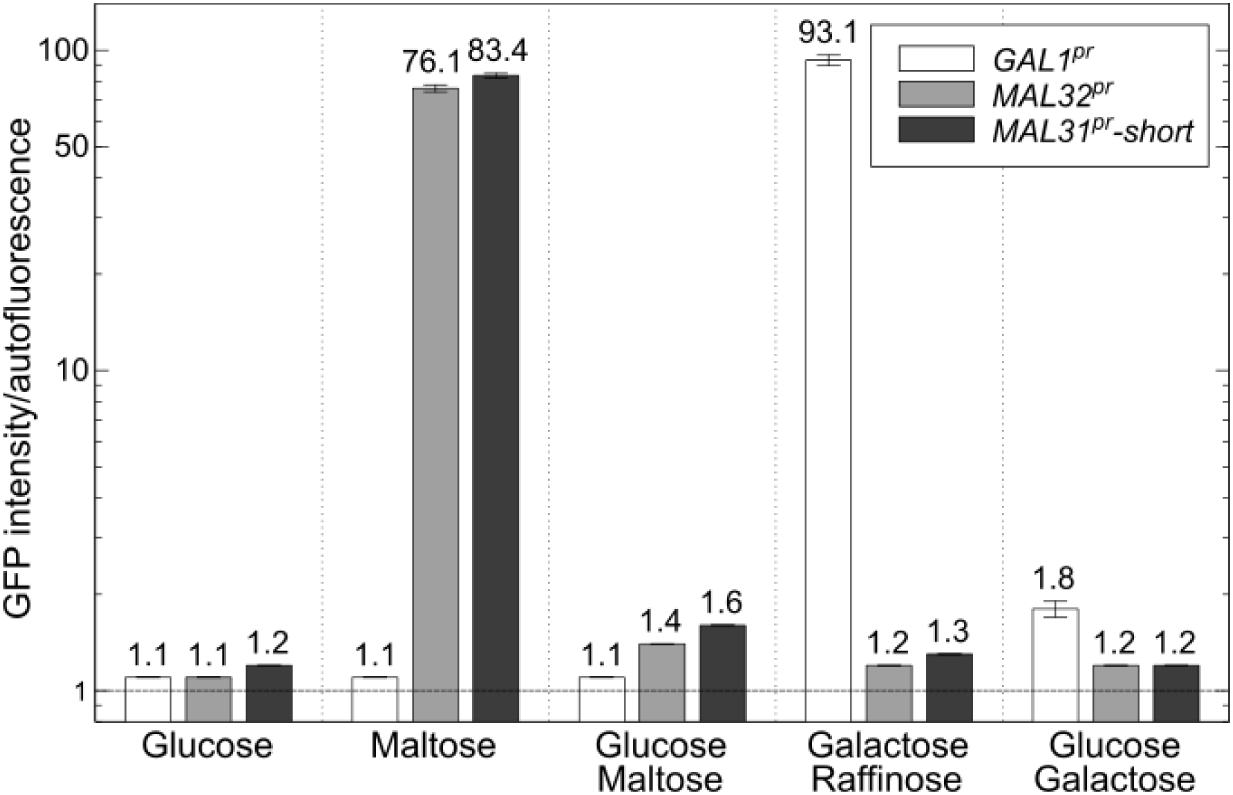
Induction and repression of *MALxy* promoters in comparison to the *GAL1* promoter. GFP fluorescence measurements using flow cytometry of strains (OD 0.5) expressing *sfGFP* driven by *GAL1^pr^* or *MALxy^pr^* from a low copy number plasmid (pRS415 (Sikorski and Hieter 1989)) as a function of different carbon sources, as indicated (2% w/v each), grown in synthetic complete media lacking leucine for > 15 hours. GFP intensities were normalized to autofluorescence of cells harboring an empty plasmid (mean ± s.d., *n* = 3 colonies per construct).

In contrast to these results obtained in liquid medium, we observed that the presence of glucose in maltose or galactose containing agar plates did not repress expression of the *MAL32^pr^* and *GAL1^pr^* constructs (data not shown). We attribute this to the fact that cell colonies on agar plates are 3D objects that receive their nutrients from the bottom. Therefore, we speculate that the cells at the top of the colony, where the fluorescence is measured using the plate reader, receive only maltose or galactose, because glucose, which is the preferred carbon source, is consumed completely by the cells underneath.

To explore the possibility to use both promoters simultaneously in experiments where orthogonal regulation of two genes is needed, we used cells harboring *MAL32^pr^* and *GAL1^pr^* fusions simultaneously, using liquid growth conditions and full induction with the corresponding carbon sources (> 15 hours of growth under inducing conditions). On maltose only the *MAL32^pr^* reporter was expressed, and on galactose only the *GAL1^pr^* reporter was expressed. In the simultaneous presence of maltose and galactose both reporter were expressed to levels reaching approximately 70% of the ones observed for ‘single sugar’ induction (Fig. 4). We analyzed this culture by flow cytometry and found that it consisted of a homogeneous population of cells where each cell expresses both reporters simultaneously (data not shown).

**Figure 4.**
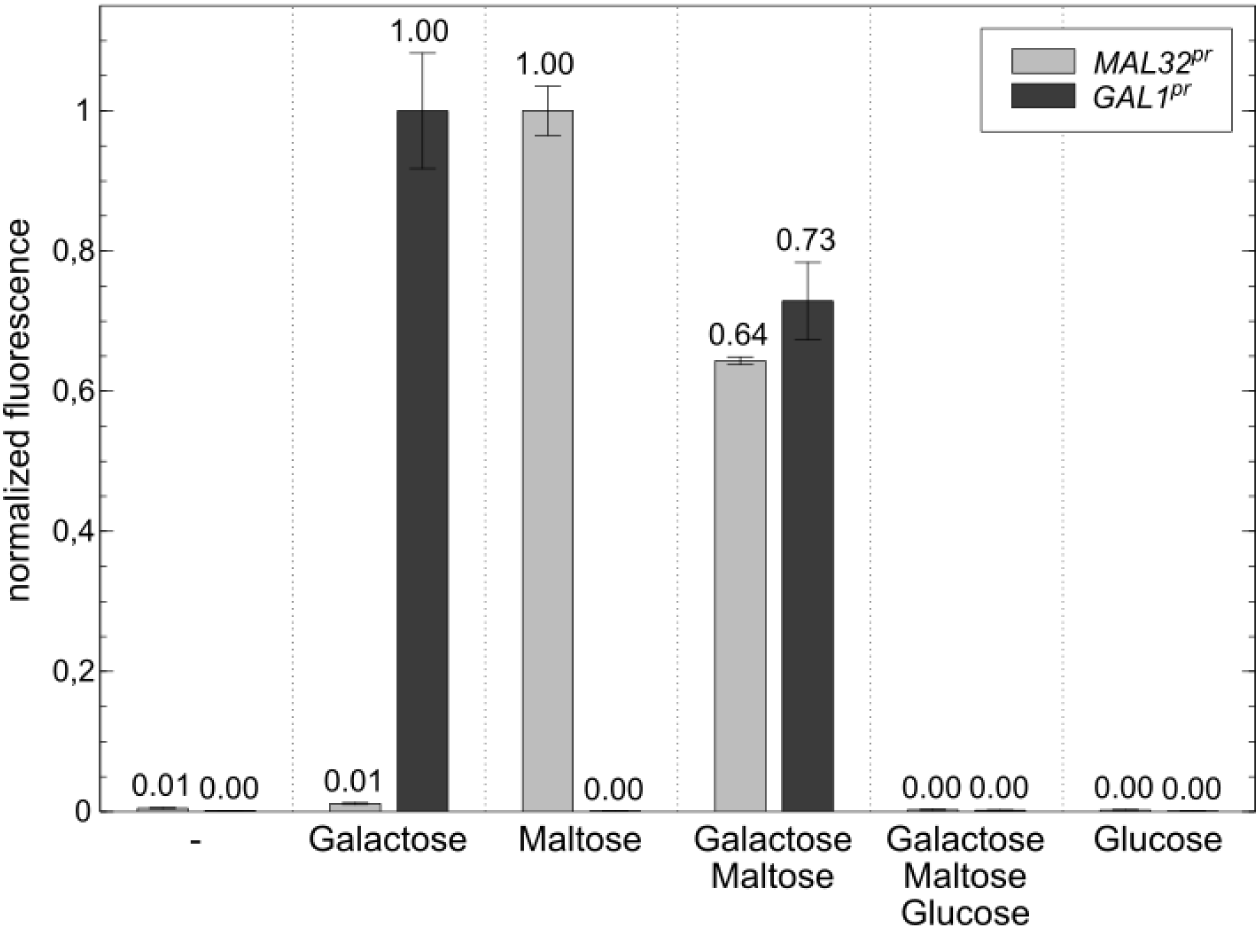
Independent regulation of *MAL32^pr^* and *GAL1^pr^*. Flow cytometry measurements of the activity of *MAL32^pr^* and *GAL1^pr^* as a function of different carbon sources as indicated (2% w/v each, 2% raffinose always included). Promoter-reporter fusions were each contained on a different plasmid in the same strain (pRS415-*MAL32^pr^-sfGFP*, pRS416-*GAL1^pr^-mCherry*). Strains were grown in synthetic complete media lacking leucine and uracil containing the indicated carbon sources for > 15 hours and measured at OD 0.5. Median autofluorescence intensities of wild type yeast colonies harboring empty plasmids were used for background correction and the measured fluorescence was normalized using values from the fully induced *MAL32^pr^* and *GAL1^pr^* reporter constructs, respectively (mean ± s.d., *n* = 3 colonies per construct).

In a last series of experiments we used flow cytometry and liquid growth conditions to compare the induction and repression dynamics of both promoters. We integrated the *GAL1^pr^* or the *MAL32^pr^* upstream of a strain expressing *NUP2-sfGFP*. We choose *NUP2* as a test case since we knew from unpublished work that overexpression or deletion does not compromise the fitness of the cell (data not shown). For *GAL1^pr^* the entire cell population showed homogeneous induction of the reporter upon galactose addition. The situation was different for *MAL32^pr^*. Here, induction did not occur uniformly in all cells simultaneously. Instead, up to 5 hours after the addition of maltose the populations still contained cells that had not (yet) induced the reporter. Only after prolonged growth in the presence of maltose for > 15 hours did all cells exhibited uniform expression (Fig.5A). Therefore *MAL32^pr^* cannot be used for short-term expression experiments where homogeneously induced cell populations are needed.

**Figure 5.**
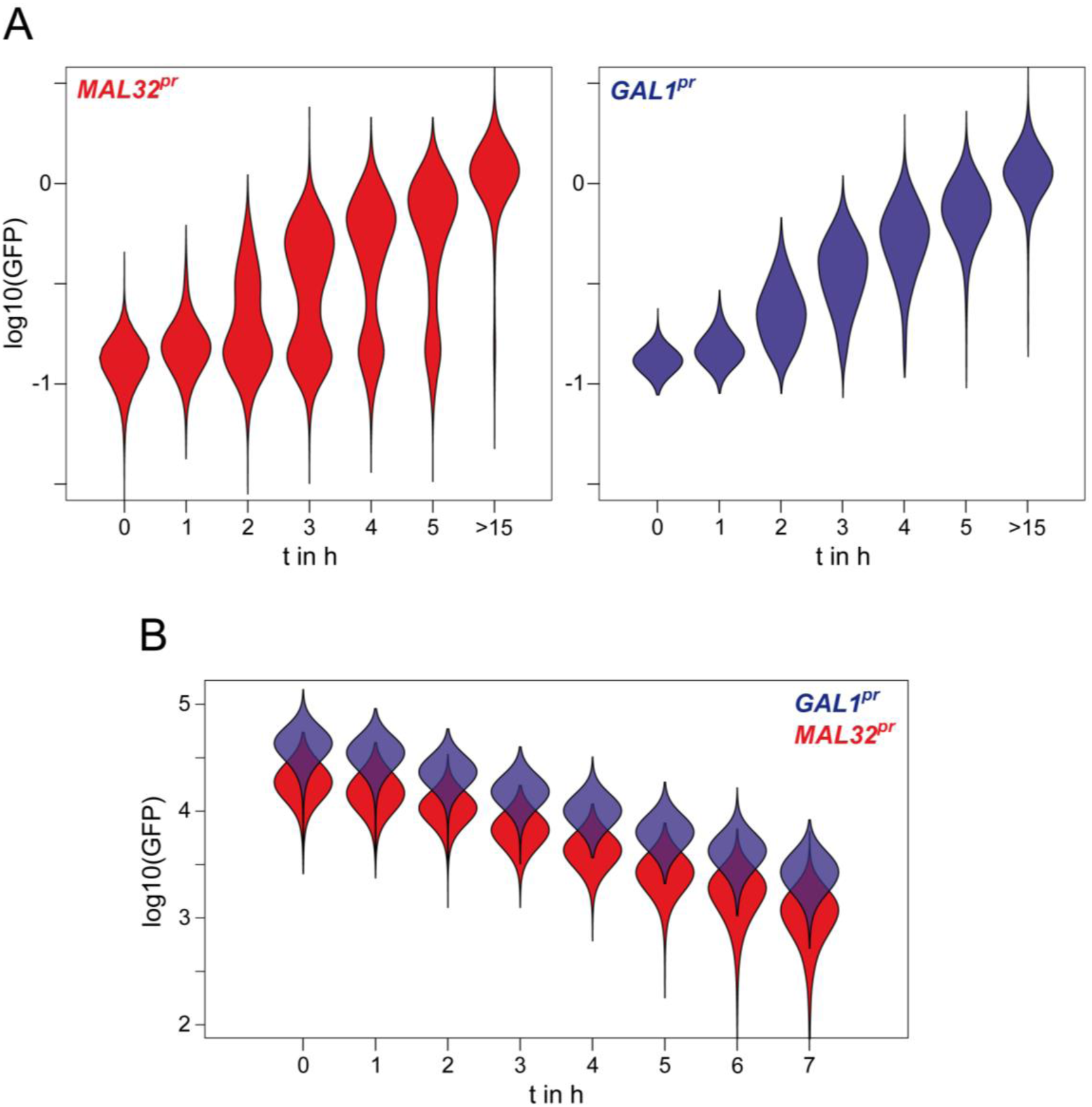
Induction and repression kinetics of *MAL32^pr^* and *GAL1^pr^*. (**A**) Induction kinetics. Time course of GFP fluorescence measurements with flow cytometry of strains expressing *NUP2*-*sfGFP* fusion driven by *GAL1^pr^* and *MAL32^pr^*. Cells were grown in synthetic complete medium containing raffinose (2% w/v) as a carbon source over night to mid log phase before induction of expression. For induction, galactose or maltose (2% w/v) was added to the diluted cultures (0.1 OD) at t = 0 min. Median autofluorescence intensity of a wild-type yeast strain was subtracted from GFP intensities and the measured values were normalized to the median value of fully induced cells. Violin plots are shown, plotted using a log scale. (**B**) Repression kinetics. Time course of GFP fluorescence measurements using flow cytometry of strains expressing *NUP2-sfGFP* driven by *GAL1^pr^* and *MAL32^pr^*. Strains were grown in synthetic complete medium containing galactose/raffinose or maltose/raffinose (each 2% w/v) as carbon sources over night to mid log phase before repression. Glucose (2% w/v) was added to the diluted cultures (0.1 OD) at t = 0 min to repress promoter activity. Median autofluorescence intensity of a wild-type yeast strain was subtracted from GFP intensities. Violin plots are shown plotted using log scale.

Glucose addition to cells on maltose or galactose medium respectively led to rapid shut down of both promoters and the cellular fluorescence decayed exponentially, indicating an immediate halt of the reporter expression and subsequent dilution of the reporter in the dividing cell population (Fig. 5B). No difference in the behavior between the *GAL1* and *MAL32* promoter was observed.

Taken together, our experiments show that *MAL32^pr^* is strongly induced by maltose in the presence of a functional *MAL*-activator and background expression from it under repressing or non-inducing conditions is minimal and comparable to *GAL1^pr^*. The *MAL32^pr^* can be used in combination with *GAL1^pr^*, provided that (i) a functional *MAL-* activator gene is present (which can be integrated into the genome or contained on a plasmid or PCR tagging cassette), (ii) induction of the *MAL32^pr^* is not time-critical, since complete induction in every cell in a culture takes up to 15 hours. While this latter property prevents the *MAL32* promoter for applications where short induction pulses are needed, it still works for experiments where longer induction periods can be accommodated (e.g. over night growth).

The diversity of *MAL* genes and the number of *MAL*-loci in different *S. cerevisiae* strains and also in other yeast species has been explored to some extent (e.g. see (Vidgren, Ruohonen and Londesborough 2005; Brown, Murray and Verstrepen 2010). However for practical reasons, e.g. when planning to use the *MAL32^pr^* for experimental work, it is only required to know whether a particular yeast strain contains a functional *MAL*-activator gene, whether it is able to grow on maltose, and whether the *MAL32^pr^* is induced. To obtain corresponding information for any yeast strain, the plasmids pMaM440 (containing a *MAL32^pr^-sfGFP*) and pMaM454 (containing a *MAL32^pr^-sfGFP* and the *MAL63* activator) can be used. In Table 1 we demonstrate this for a number of frequently used laboratory strains of diverse origin. The obtained results emphasize that laboratory strains indeed differ with respect to maltose growth and induction.

**Table 1.** *MAL*-activator availability in different wild-type yeast strain backgrounds.

| Strain | Background | pMaM440<br>(pRS415-*MAL32<sup>pr</sup>*-sfGFP) |  | pMaM454 (pRS415-*MAL63*-*MAL32<sup>pr</sup>*-sfGFP) |  |
| --- | --- | --- | --- | --- | --- |
|  |  | Growth on<br>SC + Maltose | Promoter<br>activity of<br>*MAL32<sup>pr</sup>* on<br>SC + Maltose | Growth on<br>SC + Maltose | Promoter<br>activity of<br>*MAL32<sup>pr</sup>* on<br>SC + Maltose |
| BY4741 | S288c | - | n.a. | + | + |
| SEY6210 | SEY6210 | - | n.a. | + | + |
| KN699 | W303-1A | - | n.a. | + | + |
| YPH499 | YNN216 | - | n.a. | + | + |
| YHUM216 | Sigma1278b | + | + | + | + |
| LH175 | SK1 | + | + | + | + |

n.a. – not applicable

To facilitate the use of the *MAL32^pr^* we have also constructed some tools. For N-terminal tagging of genes, tagging cassettes harboring a selection marker (*kanMX*, *hphNT1* or *natNT2*) and the *MAL32^pr^* can be used together with S1-/S4-primers (Janke *et al.* 2004) (Fig. 6B (ii), pMaM446/448/447). For induction of the promoter a yeast strain containing a functional *MAL* activator is needed (see Table 1). Since BY4741 is widely used as a laboratory strain, we have constructed a BY4741 strain with a marker free integration of the *MAL63* activator (YMaM991). For strains without a functional *MAL* activator, tagging with the *MAL32^pr^* and integration of *MAL63* can be done simultaneously together with a selection marker (Fig. 6B (i), pMaM458/456/460). For marker free integration into a strain without functional *MAL* activator a tagging cassette only containing the *MAL32^pr^* and *MAL63* can be used (Fig. 6B (iii), pMaM462). In this case selection for positive transformants needs to be done on YP + 2 % maltose + Antimycin A (3 mg/L), which enhances the selection of fermentatively growing transformants with a functional *MAL* activator over non-transformed cells that can only grow by respiration of maltose without *MAL63* (Fukuhara 2003)). One has to note that tagging efficiencies are dependent on the cassette features and their homology to the genome. For cassettes including *MAL32^pr^* or *GAL1^pr^* the tagging efficiency is about 50 %, for cassettes containing *MAL32^pr^* plus *MAL63* the efficiency goes down to 5 - 10 %. In addition to the tagging cassettes, centromeric plasmids based on pRS415 (Sikorski and Hieter 1989) harboring the *MAL32^pr^* with and without *MAL63* are available (Fig. 6A (ii)/(i), pMaM453/449).

**Figure 6.**
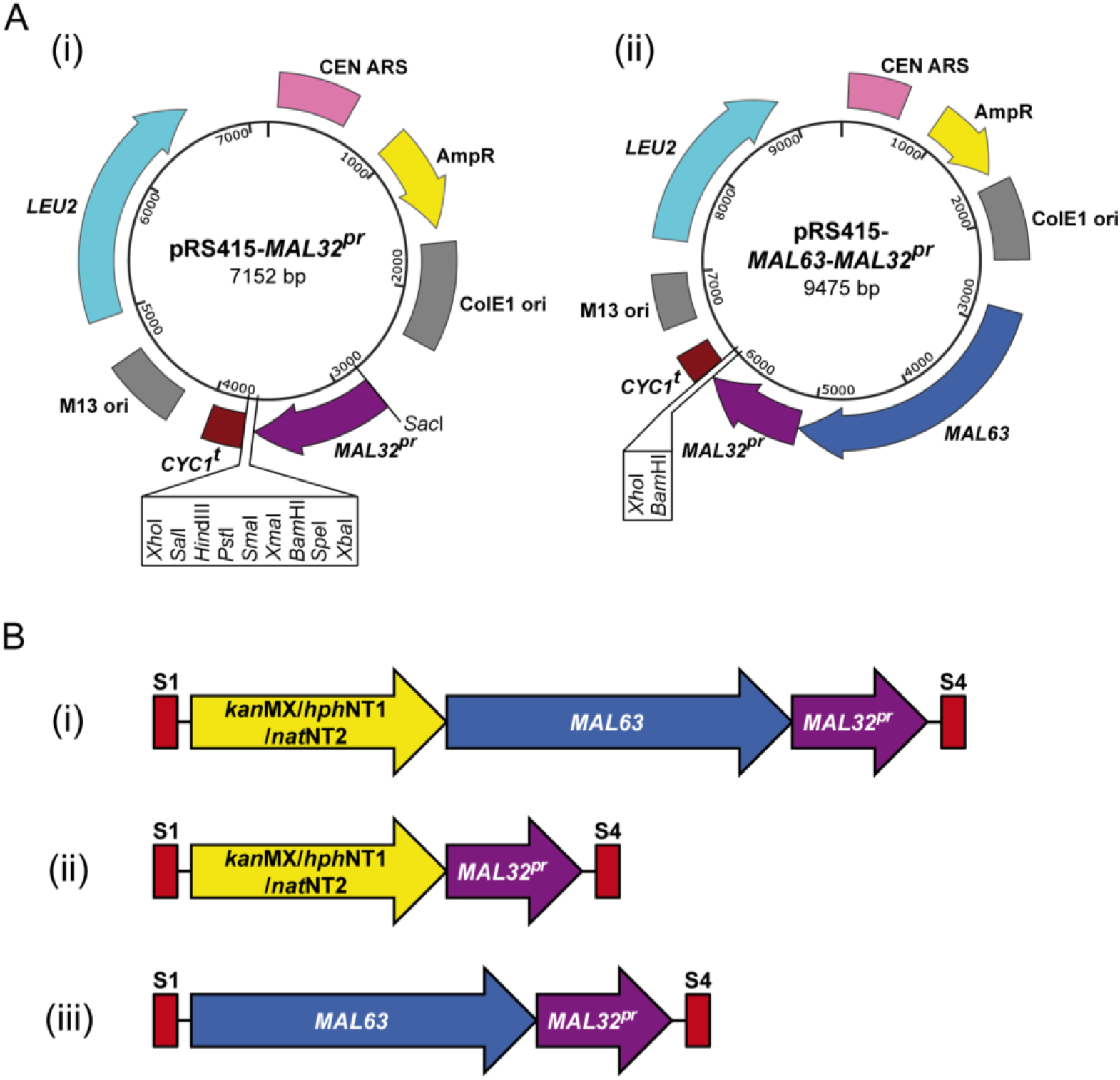
Plasmid maps. (**A**) Available yeast - *E. coli* shuttle plasmids with the *MAL32* promoter (*MAL32^pr^*), without (i) and with (ii) the activator gene *MAL63*. Sequence files can be obtained upon request. (**B**) Cassettes for PCR targeting (Maeder, Maier and Knop 2007). S1 and S4 denote PCR annealing sites for oligos commonly used for homology directed genome insertion (Janke *et al.* 2004).

i. Cassettes containing a marker as indicated, the *MAL63* activator gene and the *MAL32* promoter (for strains without functional *MAL* activator (see Table 1).
ii. Cassettes containing a marker as indicated and the *MAL32* promoter (for strains with functional *MAL* activator (see Table 1).
iii. Cassettes containing the *MAL63* activator gene and the *MAL32* promoter (for strains without functional *MAL* activator (see Table 1).

## Materials and Methods

### Yeast strains, plasmids and growth conditions

Yeast cells were grown according to standard methods (Sherman, 2002). Cultures were grown at 30°C to logarithmic phase (OD_600_ between 0.5 and 1.0 corresponding to 0.5−1=10^7^ cells/ml) unless otherwise stated. See Table 2 for a list of strains. For construction of yeast strains, standard methods were used. For chromosomal fluorescent protein reporter fusions, a one-step PCR targeting procedure was used to chromosomally introduce the fluorescent protein 3’- to the selected promoter region, while fully maintaining the integrity of the target locus, according to the method described in Huber et al. (2014) (Huber *et al.* 2014). For plasmids (Table 3), standard cloning procedures were used (Greene and Sambrook 2012). A list of primers used in this study can be found in Table 4.

**Table 2.** Yeast strains used in this study.

| Strain | Background | Genotype | Figure | Reference |
| --- | --- | --- | --- | --- |
| FY1679 | S288C | *MATa/α ura3-52/ura3-52*<br>*trp1Δ63/TRP1 leu2Δ1/LEU2*<br>*his3Δ200/HIS3 GAL2+/GAL2+* |  | (Wach *et al.* 1997) |
| ESM356-1 | FY1679 | *MATa ura3-52 leu2Δ1 his3Δ200 trp1Δ63* | 5 | Spore of FY1679 |
| BY4741 | S288C | *MATa his3Δ1 leu2Δ0 met15Δ0 ura3Δ0* | Table 1 | (Brachmann *et al.* 1998) |
| KV1042 | BY4741 | *Δmal13::hph-RM11_MAL63c9* |  | (Voordeckers *et al.* 2012) |
| YMaM991 | BY4741 | ChrXI:645199-645230Δ::RM11_MAL63c9*<br>*MAL63 integrated between genes *SIR1* and *FLO10* on Chr XI. |  | this study |
| YMaM962 | KV1042 | pRS415-MAL32<sup>pr</sup>-sfGFP (pMaM440) | 3 | this study |
| YMaM963 | KV1042 | pRS415-MAL31<sup>pr</sup>-short-sfGFP (pMaM441) | 3 | this study |
| YMaM967 | KV1042 | pRS415-GAL 1<sup>pr</sup>-sfGFP (pMaM442) | 3 | this study |
| YMaM986 | KV1042 | pRS415 pRS416 | 4 | this study |
| YMaM987 | KV1042 | pRS415-MAL32<sup>pr</sup>-sfGFP (pMaM440)<br>pRS416-GAL 1<sup>pr</sup>-mCherry (pMaM450) | 4 | this study |
| YAK947 | ESM356-1 | *NUP2::sfGFP-kanMX* |  | this study |
| YMaM992 | YAK947 | *natNT2-MAL63-MAL32<sup>pr</sup>::NUP2::sfGFP-kanMX* | 5 | this study |
| YMaM998 | YAK947 | *natNT2-GAL 1<sup>pr</sup>::NUP2::sfGFP-kanMX* | 5 | this study |
| SEY6210 | SEY6210 | *MATα leu2-3,112 ura3-52 his3-Δ200 trp1-Δ901 suc2-Δ9 lys2-801; GAL* | Table 1 | (Robinson *et al.* 1988) |
| KN699 | W303-1A | *MATa ade2-1 trp1-1 can1-100 leu2-3,112 his3-11,15 ura3-1* | Table 1 | (Thomas and Rothstein 1989) |
| YPH499 | YNN216 | *MATa ura3-52 lys2-801<sup>amber</sup> ade2-101<sup>ochre</sup> trp1Δ63 his3Δ200 leu2Δ1* | Table 1 | (Sikorski and Hieter 1989) |
| YHUM216 | Sigma1278 b | *MATa ura3-52 his3 leu2* | Table 1 | H.U. Moesch |
| LH175 | SK1 | *MATa ho::hisG lys2 ura3 leu2 his3 trp1ΔFA* | Table 1 | Linda S. Huang/Ira Herskowitz |

**Table 3.** Plasmids used in this study.

| Name | Backbone | Description | Reference |
| --- | --- | --- | --- |
| pRS415 | - | *CEN ARS LEU2* | (Sikorski and Hieter 1989) |
| pRS416 | - | *CEN ARS URA3* | (Sikorski and Hieter 1989) |
| pFA6a-*hphNT1* | - | - | (Janke *et al.* 2004) |
| pYM-N14 | - | pYM-N-*kanMX4-GPD<sup>pr</sup>* | (Janke *et al.* 2004) |
| pYM-N15 | - | pYM-N-*natNT2-GPD<sup>pr</sup>* | (Janke *et al.* 2004) |
| pYM-N23 | - | pYM-N-*natNT2-GAL 1<sup>pr</sup>* | (Janke *et al.* 2004) |
| pMaM214 | pYM-N14 | pYM-N-*hphNT1-GPD<sup>pr</sup>* | this study |
| pRS415-*MAL63* | pRS415 | pRS415-*MAL63* | this study |
| pMaM449 | pRS415 | pRS415-*MAL32<sup>pr</sup>* | this study |
| pMaM453 | pRS415 | pRS415-*MAL63-MAL32<sup>pr</sup>* | this study |
| pMaM440 | pRS415 | pRS415-*MAL32<sup>pr</sup>-sfGFP* | this study |
| pMaM441 | pRS415 | pRS415-*MAL31<sup>pr</sup>-short-sfGFP* | this study |
| pMaM454 | pRS415 | pRS415-*MAL63-MAL32<sup>pr</sup>-sfGFP* | this study |
| pMaM442 | pRS415 | pRS415-*GAL 1<sup>pr</sup>-sfGFP* | this study |
| pMaM450 | pRS416 | pRS416-*GAL1<sup>pr</sup>-mCherry* | this study |
| pMaM446 | pYM-N14 | pYM-N-*kanMX4-MAL32<sup>pr</sup>* | this study |
| pMaM447 | pYM-N15 | pYM-N-*natNT2-MAL32<sup>pr</sup>* | this study |
| pMaM448 | pMaM214 | pYM-N-*hphNT1-MAL32<sup>pr</sup>* | this study |
| pMaM458 | pYM-N14 | pYM-N-*kanMX4-MAL63-MAL32<sup>pr</sup>* | this study |
| pMaM460 | pYM-N15 | pYM-N-*natNT2-MAL63-MAL32<sup>pr</sup>* | this study |
| pMaM456 | pMaM214 | pYM-N-*hphNT1-MAL63-MAL32<sup>pr</sup>* | this study |
| pMaM462 | pMaM456 | pYM-N-*MAL63-MAL32<sup>pr</sup>* | this study |
| pMaM2 | pRS415 | pRS415-*GPD<sup>pr</sup>-sfGFP* | Ref.(Khmelins<br>kii *et al.* 2016) |
| pMaM60 | pFA6a-<br>*hphNT1* | pFA6a-*mCherry-sfGFP-hphNT1* | Ref.(Khmelins<br>kii *et al.* 2012) |

**Table 4.**
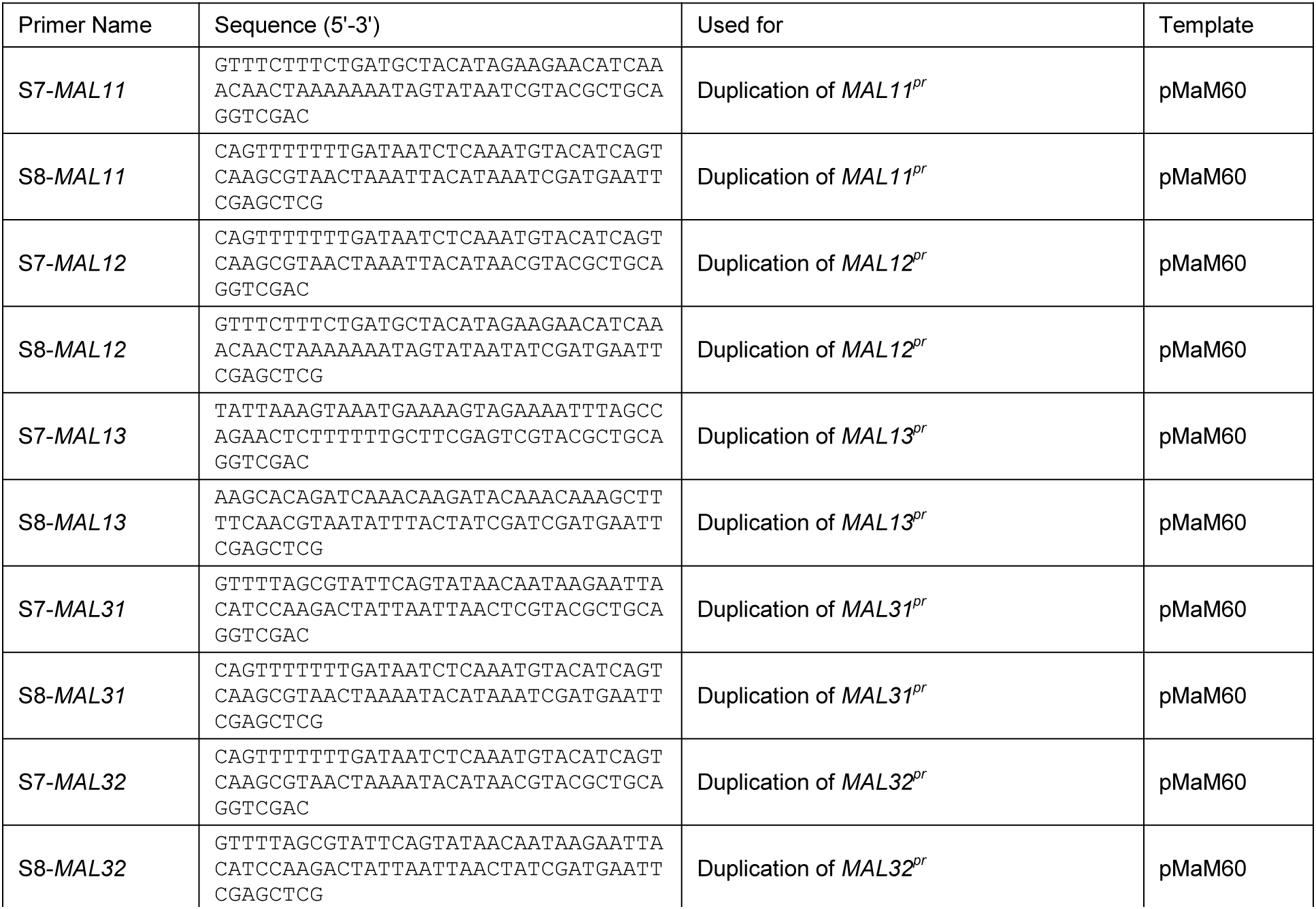
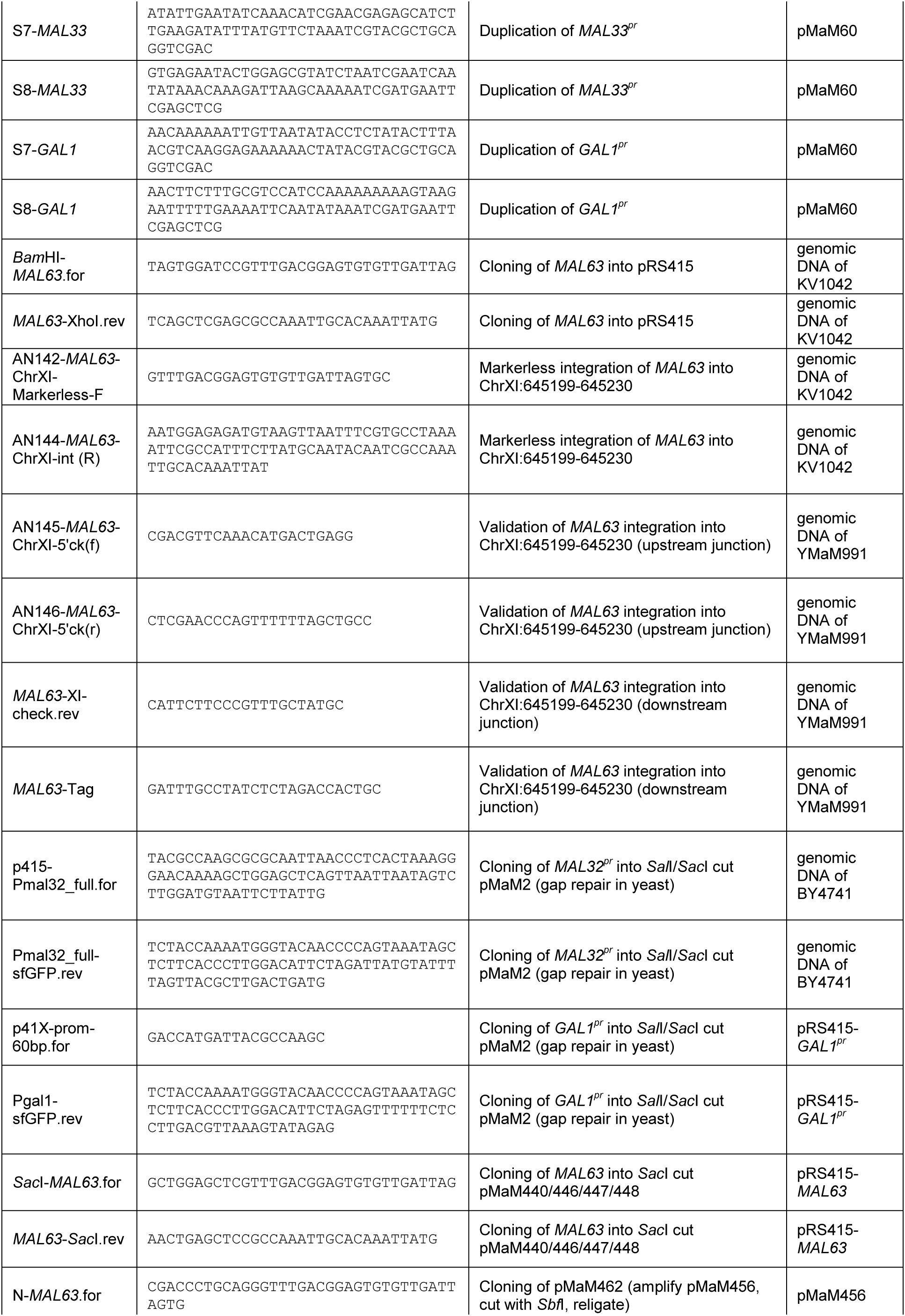
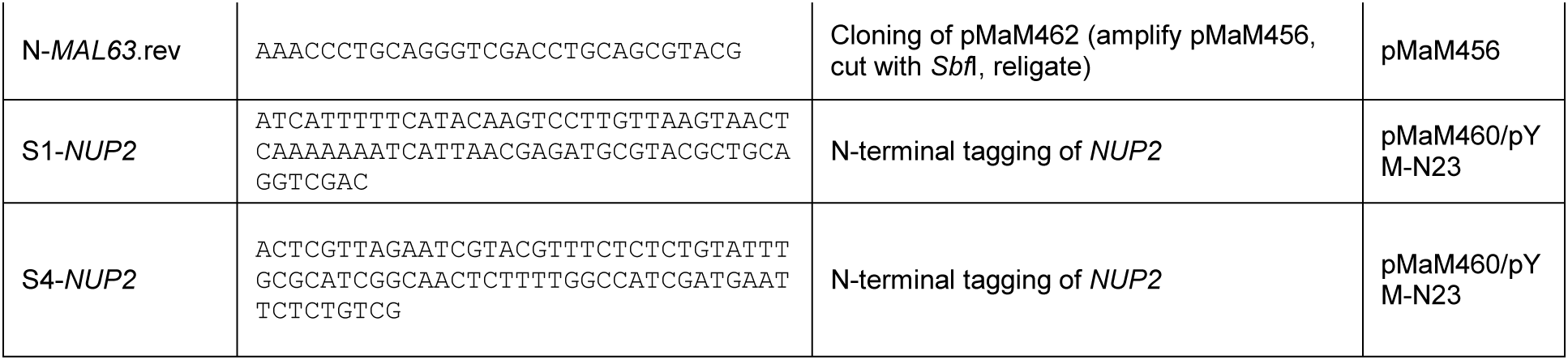
Oligonucleotides used in this study.

### Fluorescence measurements by plate reader and flow cytometry

To measure the fluorescence of colonies expressing the fluorescent protein fusions to different regulatory expression sequences, yeast colonies were pinned using a RoToR pinning robot from Singer Instruments (UK). For background subtraction of the autofluorescence of the cells we used colonies of a strain that did not express the corresponding fluorescent proteins (cells containing empty plasmids). Fluorescence was measured using a TECAN M1000 pro and appropriate settings for excitation and emission wave-lengths and gain levels for detection sensitivity. For flow cytometry a BD FACS Canto II (BD Bioscience) was used. Cells were grown to logarithmic phase (approx. 5×10^6^ cells/ml) for at least 6 hours on synthetic complete medium containing the indicated sugar (2 % w/v each).

## Acknowledgements

We thank Anton Khmelinskii for discussion and comments on the manuscript. Part of the work was funded through the DFG Grant SFB 1036.

